# *Toxoplasma* TgATG9 is critical for autophagy and long-term persistence in tissue cysts

**DOI:** 10.1101/2020.05.13.093401

**Authors:** David Smith, Geetha Kannan, Isabelle Coppens, Fengrong Wang, Hoa Mai Nguyen, Aude Cerutti, Tracey L. Schultz, Patrick A. Rimple, Manlio Di Cristina, Sébastien Besteiro, Vern B. Carruthers

**Author notes:** Corresponding authors: Vern B. Carruthers, David Smith.

## Abstract

Many of the world’s warm-blooded species are chronically infected with *Toxoplasma gondii* tissue cysts, including up to an estimated one third of the global human population. The cellular processes that permit long-term parasite persistence within the cyst are largely unknown, not only for *T. gondii* but also for related coccidian parasites that impact human and animal health. A previous study revealed an accumulation of autophagic material in the lysosome-like Vacuolar Compartment (VAC) of chronic stage bradyzoites lacking functional cathepsin L protease (TgCPL) activity. Furthermore, it was shown that TgCPL knockout bradyzoites have compromised viability, indicating the turnover of autophagic material could be necessary for bradyzoite survival. However, the extent to which autophagy itself contributes to bradyzoite development and fitness remained unknown. Herein we show that genetic ablation of *TgATG9* substantially reduces canonical autophagy and compromises bradyzoite viability. Transmission electron microscopy revealed structural abnormalities occurring in Δ*atg9* bradyzoites, including disorganization of the inner membrane complex and plasma membrane, the occurrence of multiple nuclei within a single bradyzoite cell, as well as various anomalies associated with the VAC. TgATG9-deficient bradyzoites accumulated significantly less undigested material in the VAC upon inhibition of TgCPL activity, suggesting that autophagy contributes material to the VAC for degradation. Intriguingly, abnormal mitochondria networks were observed in TgATG9-deficient bradyzoites. They were thin and elongated and often adopted a horseshoe conformation. Some abnormal mitochondrial structures were found to contain numerous different cytoplasmic components and organelles. Bradyzoite fitness was found to be drastically compromised, both *in vitro* and in mice, with very few brain cysts identified in mice 5 weeks post-infection. Taken together, our data suggests that TgATG9, and by extension autophagy, is critical for cellular homeostasis in bradyzoites and is necessary for long-term persistence within the cyst of this coccidian parasite.

## INTRODUCTION

The subclass Coccidiasina (coccidia hereafter) includes several notable parasites that are important to public and animal health (1). A unifying feature of Coccidia is their ability to form persistent cysts. All coccidian parasites generate environmentally resilient oocysts that persist in soil or water for fecal-oral transmission (2,3). Also, some Coccidia (those of the family Sarcocystidae, meaning “flesh cyst”) including *Toxoplasma gondii* form long-lived tissue cysts that confer transmission via carnivorism (3). Coccidia are further classified within the phylum Apicomplexa along with haemosporidians including *Plasmodium* spp. and *Babesia* spp., the agents of malaria and babesiosis, respectively (4). Because *T. gondii* is genetically tractable, can be grown continuously in culture, and can be injected into mice for a reliable model of the disease, this parasite has become an attractive model system for other related parasites. *T. gondii* is also recognized as an important public health pathogen, acquired through the ingestion of tissue cysts in contaminated meat, or water and food contaminated with oocysts (5). Infection with this parasite can have various serious negative health consequences (6). If acquired early during pregnancy, the parasite can traverse the placenta and infect the developing fetus, which can result in miscarriage and neurological pathologies (7). Infection can be fatal in immunocompromised individuals, for example AIDS patients, organ transplant recipients or patients undergoing chemotherapy (6). There have also been reports of emerging *T. gondii* strains causing complications and death in otherwise healthy individuals (7,8).

Within an intermediate host, sarcocystidian parasites exist in both an acute stage (tachyzoite) and a chronic stage tissue cyst (bradyzoite). During the tachyzoite phase, *T. gondii* parasites replicate rapidly within a parasitophorous vacuole (PV). Throughout the acute stage of infection, nutrients are trafficked from the host cytosol across the PV membrane (PVM) (9,10). The removal of amino acids from culture media *in vitro* triggers tachyzoite to bradyzoite differentiation (11,12). In an *in vivo* setting within an immunocompetent intermediate host, an effective host response involves CD8+ T cell recognition of host cells infected with *T. gondii* and production of interferon gamma (IFN-γ) (13,14). In turn, this results in the restriction of amino acids, including arginine and tryptophan, in cells harboring the parasite (15–17). As parasites continue to draw such resources, the infected host cell might become an increasingly nutrient-limited environment. Accordingly, it has been suggested that constrained access to essential nutrients is a major factor driving tachyzoite to bradyzoite differentiation *in vivo* (17). However, this also raises the question as to how *T. gondii* tissue cysts containing bradyzoites are able to persist within the host with potentially limited access to host material for sustenance. Once the cyst is fully developed, the extent to which mechanisms for trafficking host material across the PVM in tachyzoites can still take place across the cyst wall remains to be determined.

Macroautophagy (hereafter autophagy, for “self-eating”) is an intracellular catabolic process that facilitates the encapsulation, trafficking and degradation of endogenous proteins and organelles (18,19). This process begins with the activation of an initiation complex comprised of several proteins including the protein kinase ATG1. ATG1 activity catalyzes several downstream events including the assembly of a complex comprising the lipid transfer protein ATG2, the proppin family protein ATG18 and the only integral membrane protein in the pathway ATG9. The ATG2/18/9 complex facilitates the development of a double membraned phagophore that elongates while capturing cytoplasmic material. During the process of elongation, the small ubiquitin-like protein ATG8 accumulates on the developing phagophore via lipid conjugation to phosphatidylethanolamine. The phagophore closes to generate a double membrane vesicle, the autophagosome. Fusion of the autophagosome with a lytic compartment such as the lysosome (vertebrates) or the vacuole (yeast and apicomplexan parasites) creates an autolysosome within which cytoplasmic material is degraded by cathepsin proteases and other hydrolytic enzymes. The entire process of development, closure and fusion occurs quite rapidly, often within 3-10 min (20,21).

The turnover of autophagic material via the digestive organelle can serve a number of purposes that all contribute towards cellular homeostasis and promote cell survival. These include: recycling nutrients as part of a starvation response, removal of damaged material from the cytoplasm, turnover of proteins and organelles during developmental changes and the removal of pathogens (18). The autophagy machinery is widely conserved among eukaryotes. However, whether or not all Apicomplexa are able to perform canonical autophagy is still a matter of debate, even if there are clues *T. gondii* has a functional pathway (22). Interestingly, several *Plasmodium* and *Toxoplasma* autophagy-related proteins are also involved in a non-canonical function related to the segregation of the apicoplast (23–26), a non-photosynthetic plastid shared by most members of the phylum.

It has previously been shown that while *T. gondii* cathepsin L (TgCPL) is dispensable in tachyzoites (9,27), inhibition or genetic ablation of this protease in bradyzoites results in parasite death (27,28). TgCPL is a cysteine protease that resides within a Plant-Like Vacuole (PLV)/Vacuolar Compartment (VAC, hereafter) in *T. gondii*, wherein it is a major enzyme required for the degradation of proteinaceous material (9,10,27,28). Interestingly, we previously demonstrated the accumulation of autophagic material and organellar remnants within the VAC in parasites lacking active TgCPL (28). Collectively, these findings suggested the accumulation of undigested autophagic material within the VAC in bradyzoites was somehow contributing to parasite death, although precisely why parasites were dying remained unknown (28).

In the present study, we set out to explain why TgCPL-deficient bradyzoites fail to survive and to determine the extent to which autophagy itself directly contributes to chronic persistence of *T. gondii* as a model sarcocystidian and coccidian parasite.

## RESULTS

### Conservation of canonical autophagic proteins in coccidian parasites

Previous analyses of the autophagy machinery in *Toxoplasma gondii* have shown a reasonably high degree of evolutionary conservation compared with other apicomplexan parasites (22,29). Here, we sought to determine the relative conservation of *T. gondii* autophagy genes among related apicomplexans, particularly cyst-forming parasites, as well as related but non-parasitic aveolates. Expectedly, we found that autophagy-related genes in the closely related sarcocystidian parasites *Hammondia hammondi* and *Neospora caninum* are most conserved with those of *T. gondii* (**Figure 1A**). Intriguingly, while some genes associated with canonical autophagy (e.g., ATG2 and ATG9) are also conserved in several coccidian parasites, they were not identified in the non-coccidian apicomplexan parasites *Cryptosporidium parvum, Plasmodium falciparum* and *Babesia spp*. Furthermore, autophagy-related genes tended to be more conserved between *T. gondii* and the non-parasitic aveolates *Chromera velia* and *Vitrella brassicaformis* (the closest known autotrophic organisms to the Apicomplexa) than between *T. gondii* and other non-coccidian Apicomplexa. However, matches to the autophagy-related *T. gondii* genes that are not only involved in canonical autophagy but also in maintenance of the apicoplast (which is found in all apicomplexans except *Cryptosporidium* spp.), namely ATG3, ATG7, ATG8, Prop2, VPS15 and VPS34, were found across all the protists considered in this analysis (except in *Babesia* spp., which only had ATG7, ATG8 and VPS34). Taken together this information indicates a potential role for canonical autophagy in sarcocystidia and possibly some other coccidia versus a more universal non-canonical function for autophagy-related proteins in maintenance of the apicoplast.

**Figure 1.**
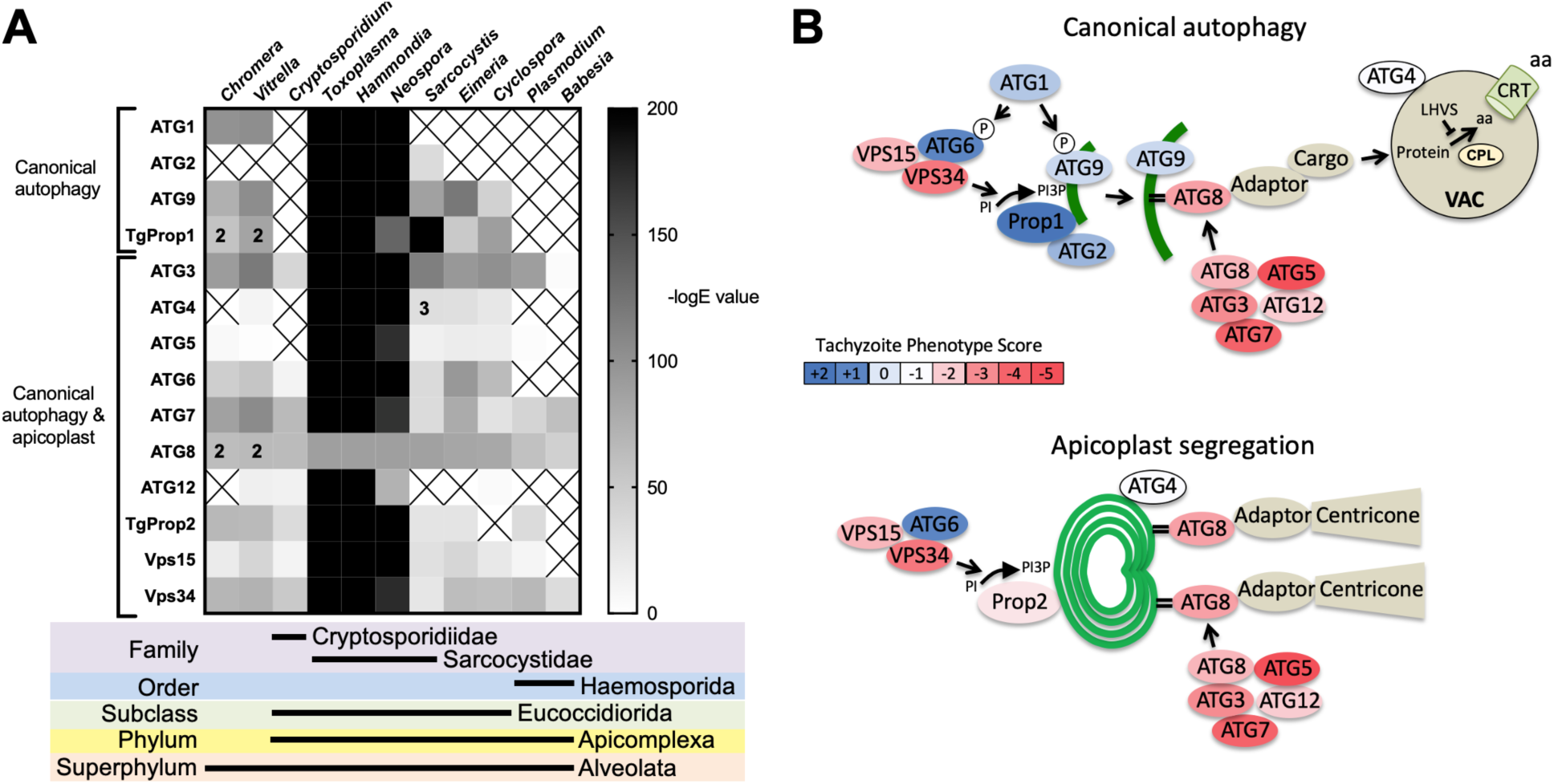
Phylogenetic conservation and adaptation of autophagy-related proteins in alveolates. (A) Reciprocal BLAST hits to *T. gondii* autophagy-related proteins detected in different alveolates. The inverse logarithm of the E value (-LogE) from each hit with a corresponding *T. gondii* protein is plotted in a heat map to indicate the extent to which *T. gondii* autophagy-related proteins are conserved among other alveolates. A higher -LogE value indicates a closer match to *T. gondii* orthologues. The number of orthologs is shown in the corresponding cell if more than one ortholog was identified. Boxes with a cross indicate no matching orthologue was identified. (B) Working models of canonical autophagy and apicoplast segregation mediated by a subset of autophagy-related proteins. Autophagy-related proteins are color coded according to phenotype scores indicated in the scale. Autophagy proteins adapted for apicoplast maintenance tend to have negative phenotype scores in tachyzoites, based on a genome-wide CRISPR/Cas9 screen in *T. gondii* (30), indicating they are essential for parasite survival in the acute stage of infection. Autophagy-related proteins that have a role in canonical autophagy and have not been adapted for apicoplast maintenance tend to have neutral or positive phenotype scores, indicating canonical autophagy is a non-essential cellular pathway in acute stage parasites.

To gain a sense of which components of the autophagy pathway in *T. gondii* play critical roles for the parasite, we mapped such constituents to a working model of autophagy and color coded them according to their phenotype scores from a recent genome-wide CRISPR-Cas9 screen (30) (**Figure 1B**). The phenotype score reflects the relative fitness cost associated with loss of each gene during *in vitro* replication of tachyzoites. A strongly negative phenotype score indicates essentiality, whereas a neutral or positive phenotype score suggests dispensability, at least during acute stage replication in culture. Interestingly, proteins that have been implicated in segregation of the apicoplast, including TgATG8 and its lipid conjugation machinery (e.g., TgATG3, TgATG5, TgATG7 and TgATG12), together with most proteins in the VPS34/PI3P kinase complex, have strongly negative phenotype scores. Conversely, proteins that are theoretically (TgATG1 and TgATG2) or empirically (TgProp1 (23) and TgATG9 (31)) linked exclusively to canonical autophagy have neutral or positive phenotype scores (**Figure 1B**). This is consistent with canonical autophagy playing a non-essential role in cultured tachyzoites. Nevertheless, that proteins involved exclusively in canonical autophagy have been retained in cyst-forming parasites suggests that their biological significance could manifest in a different stage of the infection.

### TgATG8 is required for autophagy and viability of T. gondii bradyzoites

We previously demonstrated that autophagic material and lipidated TgATG8 accumulate in TgCPL deficient bradyzoites due to lower protein turnover in the VAC (28). Although this suggested that bradyzoites undergo autophagy, the evidence was indirect. As an initial strategy to directly test for autophagy in bradyzoites, we targeted TgATG8, which is necessary for apicoplast segregation in tachyzoites (24) and presumably also required for canonical autophagy, as in other systems. To achieve selective loss of expression in bradyzoites, we used CRISPR/Cas9 to insert a tachyzoite stage specific SAG1 promoter upstream of the endogenous *TgATG8* gene in the Prugniaud (Pru, hereafter referred to as wild-type, WT) strain, thus generating S/ATG8 transgenic parasites (**Figure 2A-B**). At the same time, we also added an N-terminal GFP tag to TgATG8. S/ATG8 tachyzoites showed GFP-TgATG8 associated with the apicoplast (**Figure 2C**), which appeared to segregate normally, consistent with GFP-TgATG8 expression being sufficient to support its critical role in maintaining this organelle (24). Upon *in vitro* differentiation, expression of GFP-TgATG8 decreased to below the limits of detection by fluorescence microscopy and western blotting (**Figure 2C-D**). To assess the role of TgATG8 in bradyzoite autophagy, we converted S/ATG8 to bradyzoites for 7 days and looked for accumulation of dark puncta (indicating accumulation of undigested material in the VAC; (28)) and CytoID-positive structures (autophagic material in the VAC; (28)) following treatment with the cathepsin L inhibitor LHVS. Whereas 24 hours of LHVS treatment resulted in the expected accumulation of dark puncta and CytoID-positive structures in WT bradyzoites, such structures were lacking in S/ATG8 bradyzoites (**Figure 2E**), hinting autophagy is indeed affected in this mutant. Dark puncta and CytoID-positive structures were even more pronounced in WT bradyzoites following 7 days of continual LHVS treatment. Some small puncta and CytoID signal were present in S/ATG8 bradyzoites after 7 days of LHVS treatment, suggesting a measure of residual autophagy in the nominal absence of TgATG8 (**Figure 2E**). No viable S/ATG8 parasites were recovered from differentiated cultures upon isolating bradyzoites and applying them to fresh monolayers for plaque formation (**Figure 2F**). Although these experiments do not distinguish whether loss of viability is due to TgATG8’s role in canonical autophagy or apicoplast segregation, the findings provide an initial indication of an active autophagy pathway in bradyzoites.

**Figure 2.**
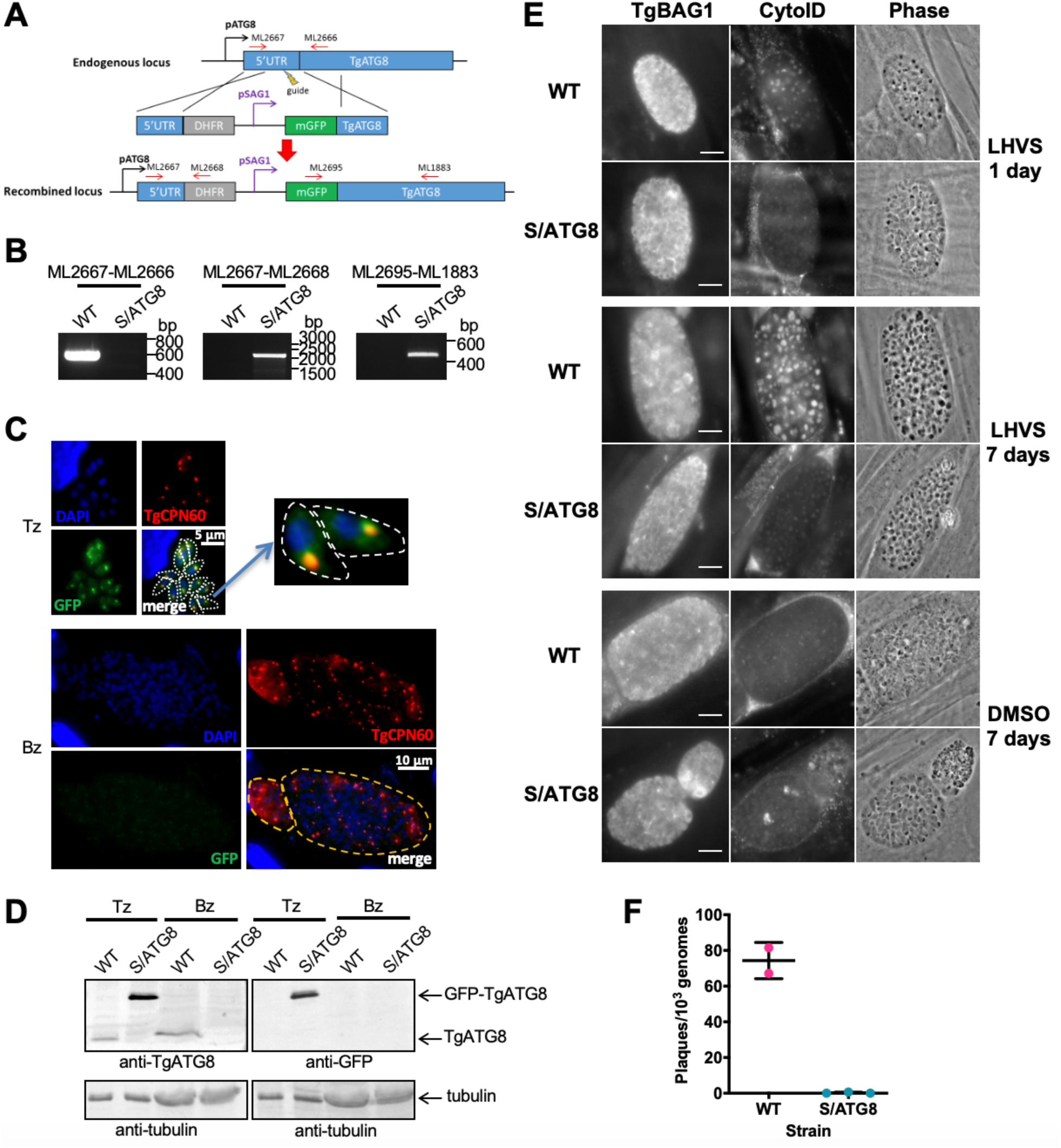
TgATG8 is required for efficient autophagosome production in *T. gondii* bradyzoites. (A) Stage-specific expression and fluorescent tagging of TgATG8 was achieved by insertion of the tachyzoite-specific *TgSAG1* promoter and a GFP coding sequence immediately upstream of the *TgATG8* gene. Site-specific insertion of the DHFR-pSAG1-mGFP repair template was performed by homology-directed repair. (B) PCR analysis confirmed the repair template had been inserted at the correct locus to generate a *T. gondii* strain in which *TgATG8* expression was driven by the *TgSAG1* promoter (*S/ATG8*). Stage-specific expression of GFP-TgATG8 in tachyzoites (Tz) but not bradyzoites (Bz) was confirmed by immunofluorescence microscopy (C) and western blotting (D). GFP-TgATG8 signal overlapped with the apicoplast marker TgCPN60, indicating correct localization to the apicoplast. Dashed lines outline individual parasites in the tachyzoite images and the limits of cysts on the bradyzoite images. (E) Fluorescent staining of *in vitro* differentiated cysts with the bradyzoite specific marker TgBAG1 and the autolysosome detection reagent CytoID. Phase contrast images indicate the development of dark puncta in LHVS treated bradyzoites. Scale bar, 10 µm. (F). Viability of WT and S/ATG8 bradyzoites isolated after 28 days of differentiation, as indicated by their application to fresh monolayers for plaque formation.

### Autophagy is a source of material for degradation in the VAC during chronic infection

To exclusively assess the role of canonical autophagy in *T. gondii* bradyzoites, it was necessary to focus on an autophagy-related protein that is required for the formation of autophagosomes but is not necessary for apicoplast maintenance. Working with a non-cystogenic strain, we previously showed the autophagy-related protein TgATG9 was dispensable in cultured tachyzoites and that TgATG9 ablation had no observable impact on apicoplast homeostasis (31). However, TgATG9-deficient parasites showed reduced protein turnover as extracellular tachyzoites, implying a role for TgATG9 in autophagy (31). Therefore, we knocked out TgATG9 in a PruΔ*hxg* background strain (also referred to as wild-type, WT, hereafter) to generate a cystogenic Δ*atg9* strain (**Figure S1A-B**). Genetic complementation was achieved by expressing *TgATG9* from its cognate promoter and 5’ and 3’ untranslated regions together with 3 copies of a C-terminal HA epitope tag, thus creating Δ*atg9ATG9*. Δ*atg9ATG9* tachyzoites expressed *TgATG9* transcript at levels comparable to those of WT based on qRT-PCR (**Figure S1C**), and expression of the TgATG9-3xHA was confirmed by immunofluorescence microscopy and western blotting (**Figure S1D-E**).

**Figure S1.**
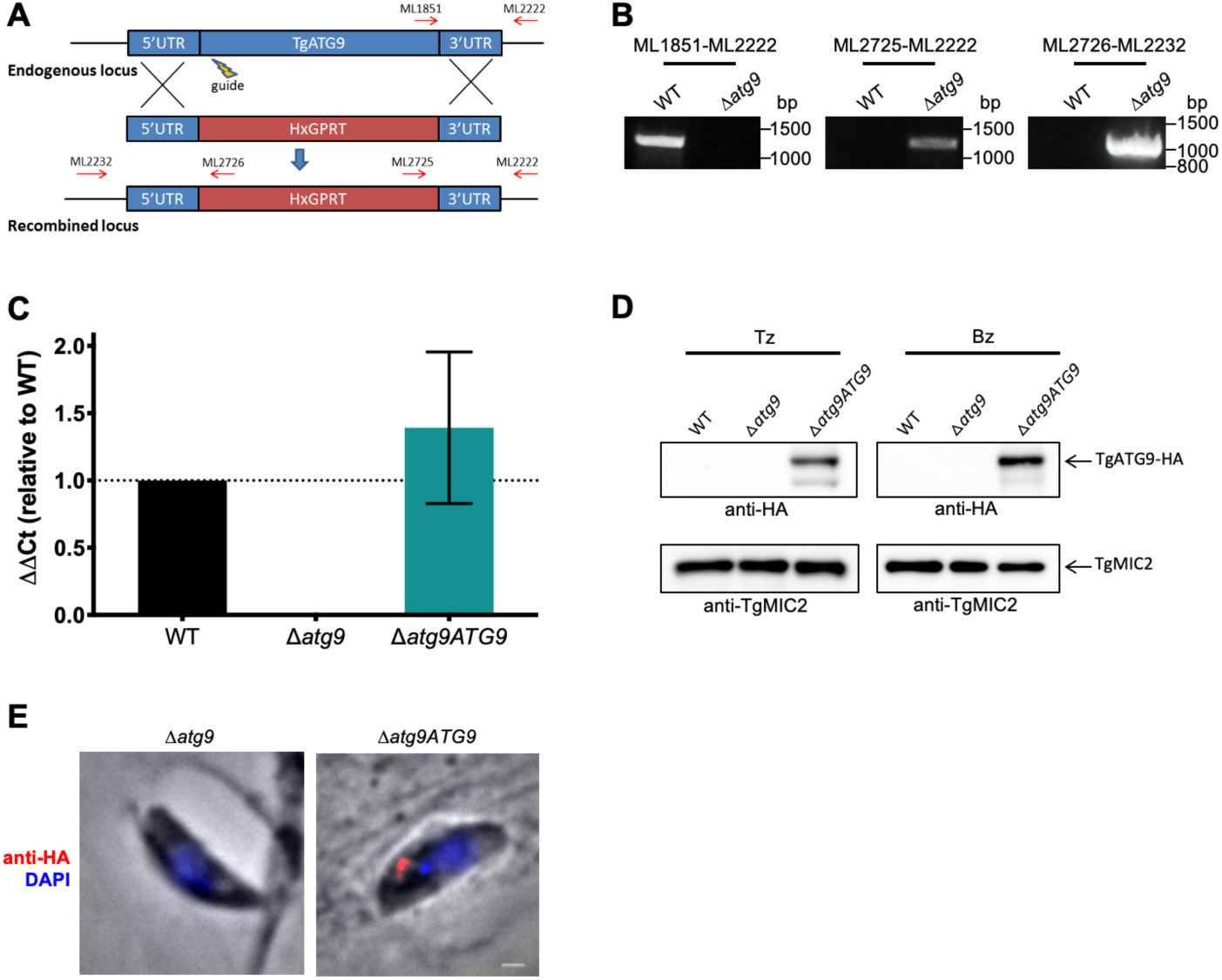
Targeted deletion and genetic complementation of TgATG9. (A) The *TgATG9* gene was replaced with an HXGPRT selectable marker by homology directed repair facilitated by CRISPR-Cas9 double strand break directed via a single sgRNA (guide). Primers used to validate correct integration are indicated as red arrows. (B) PCR validation of Δ*atg9* parasites using the primers indicated in panel A. (C) qRT-PCR analysis showing relative transcript levels of *TgATG9* in WT, Δ*atg9* and Δ*atg9ATG9* strains. (D) Western blots of tachyzoites (Tz) and bradyzoites (Bz) showing re-expression of TgATG9-HA in both stages of Δ*atg9ATG9* parasites detected with anti-HA antibodies. The same blot was stripped and reprobed with anti-TgMIC2 as a loading control. (E) IFA showing re-expression of TgATG9-HA in a newly invaded tachyzoite. Scale bar, 100 nm.

Using Δ*atg9* and Δ*atg9ATG9* transgenic strains, we next sought to confirm whether canonical autophagy is a source of material for degradation in the bradyzoite VAC. Bradyzoites were differentiated for 7 days *in vitro* and treated with either DMSO (vehicle control) or LHVS for 24 hours under differentiation conditions. Dark puncta were visible in LHVS-treated WT and Δ*atg9ATG9* bradyzoites but were less prominent in Δ*atg9* bradyzoites (**Figure 3A**). Although a significant increase in puncta size was seen in Δ*atg9* bradyzoites treated with LHVS compared to the same strain treated with DMSO, this increase was ∼64% lower than that for LHVS-treated WT and ∼73% lower than that for Δ*atg9ATG9* bradyzoites (**Figure 3B**). Also, whereas WT and Δ*atg9ATG9* bradyzoites displayed CytoID-positive structures following LHVS treatment, such structures were much less prominent in Δ*atg9* bradyzoites (**Figure 3C**). Quantification revealed that LHVS-treated Δ*atg9* bradyzoites have smaller and, importantly, fewer CytoID positive structures than those of WT and Δ*atg9ATG9* bradyzoites (**Figure 3D-E**). Taken together, these findings establish that expression of TgATG9 is necessary for efficient delivery of autophagic material to the VAC in chronic stage *T. gondii*, thus suggesting a central role for TgATG9 in canonical autophagy.

**Figure 3.**
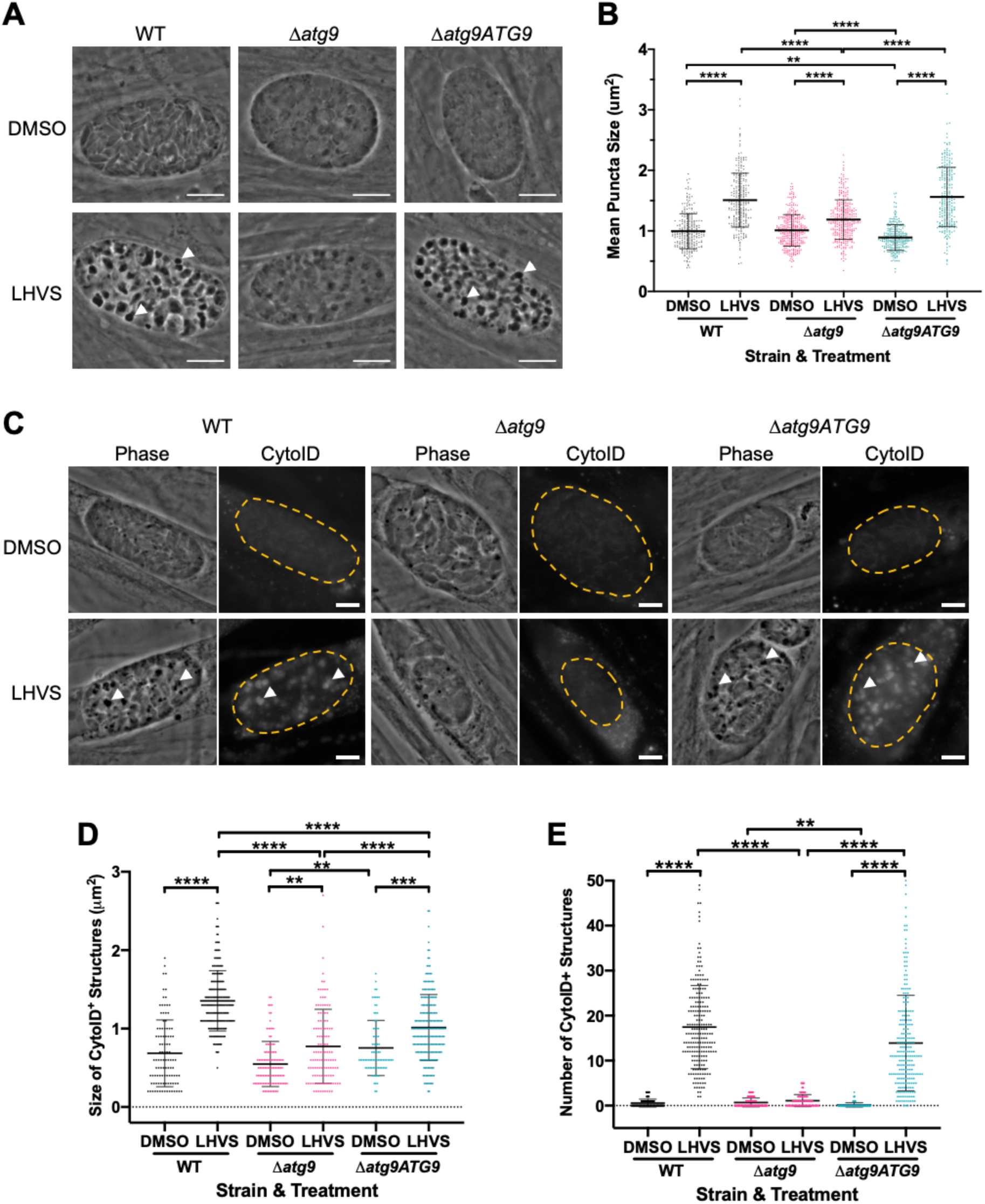
TgATG9 is required for efficient autophagosome production in *T. gondii* bradyzoites. (A) Phase contrast images showing that the development of dark puncta in *in vitro* differentiated bradyzoites following LHVS treatment is dependent on the expression of TgATG9. Arrowheads indicate a subset of dark puncta. Scale bar, 10 µm. (B) Quantification of dark puncta size. Each dot represents the mean size of puncta within one cyst. Data are merged from 6 biological replicates, with a minimum of 22 cysts analyzed per sample type, per biological replicate and a minimum total of 242 cysts analyzed per sample type across all biological replicates. (C) CytoID staining of autolysosomes in *in vitro* differentiated bradyzoite cysts. Arrowheads indicate a subset of dark puncta and the corresponding CytoID positive structures. Scale bar, 5 µm. Quantification of the size (D) and number (E) of CytoID-positive structures in *in vitro* differentiated bradyzoites. Each dot represents the average puncta measurement within a single cyst. Data are merged from 3 biological replicates, with a minimum total of 81 cysts analyzed per sample type across 3 biological replicates. For panels B, D and E, bars represent mean ± S.D. Statistical comparisons were done using a Kruskal-Wallis test with Dunn’s multiple comparisons. Statistical significance is indicated as follows: **, p< 0.01; ***, p< 0.001; ****, p<0.0001. Non-significant differences are not indicated. Statistical analysis is only shown for comparisons that have one variable i.e., different strain or different treatment.

### Canonical autophagy is required for normal bradyzoite morphology and cell division

To initially determine whether removal of the *TgATG9* gene had an effect on cyst development and bradyzoite morphology, we measured cyst size based on staining the cyst wall with *Dolichos* lectin and assessed the overall appearance of bradyzoites with the peripheral marker TgIMC1. Interestingly, we found that Δ*atg9* cysts are moderately larger than WT or Δ*atg9ATG9* cysts when measured after 7 days of differentiation (**Figure 4A**). Individual Δ*atg9* bradyzoites appeared to be misshapen and bloated within their cysts, suggesting a possible basis for cyst enlargement (**Figure 4B**). When viewed by transmission electron microscopy (TEM), WT bradyzoites showed normal ultrastructural features and typical formation of daughter cells within mother bradyzoites (**Figure 4Ca,Da**). By contrast, Δ*atg9* bradyzoites were often deformed, vacuolized and showed abnormal ultrastructural features including an enlarged nuclear envelope (**Figure 4Cb**). Some Δ*atg9* bradyzoites also showed aberrant formation or budding of daughter cells, which manifested as enlarged parasites containing multiple daughter nuclei (**Figure 4Db,Dc**). This multinucleated phenotype was found in 26% (61/235) of Δ*atg9* bradyzoites but less than 1% (2/144) of WT bradyzoites (**Table 1**). Together, these findings suggest that TgATG9 is necessary for the overall fitness of bradyzoites and its loss compromises normal bradyzoite morphology and cell division.

**Table 1.**
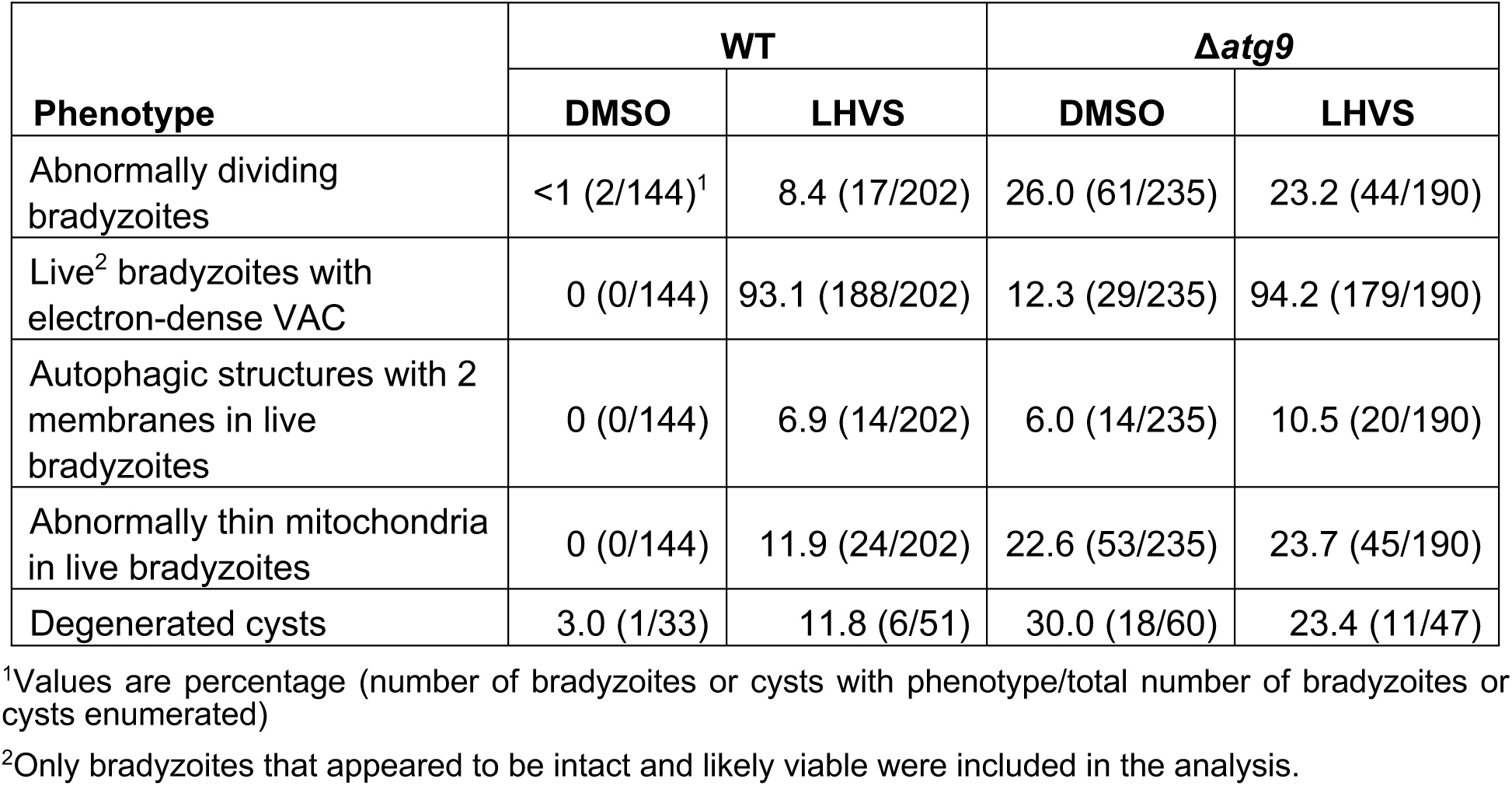
Phenotypes observed by TEM.

| Phenotype | WT |  | *Δatg9* |  |
| --- | --- | --- | --- | --- |
|  | DMSO | LHVS | DMSO | LHVS |
| Abnormally dividing bradyzoites | <1 (2/144)<sup>1</sup> | 8.4 (17/202) | 26.0 (61/235) | 23.2 (44/190) |
| Live<sup>2</sup> bradyzoites with electron-dense VAC | 0 (0/144) | 93.1 (188/202) | 12.3 (29/235) | 94.2 (179/190) |
| Autophagic structures with 2 membranes in live bradyzoites | 0 (0/144) | 6.9 (14/202) | 6.0 (14/235) | 10.5 (20/190) |
| Abnormally thin mitochondria in live bradyzoites | 0 (0/144) | 11.9 (24/202) | 22.6 (53/235) | 23.7 (45/190) |
| Degenerated cysts | 3.0 (1/33) | 11.8 (6/51) | 30.0 (18/60) | 23.4 (11/47) |

<sup>1</sup>Values are percentage (number of bradyzoites or cysts with phenotype/total number of bradyzoites or cysts enumerated)
<sup>2</sup>Only bradyzoites that appeared to be intact and likely viable were included in the analysis.

**Figure 4.**
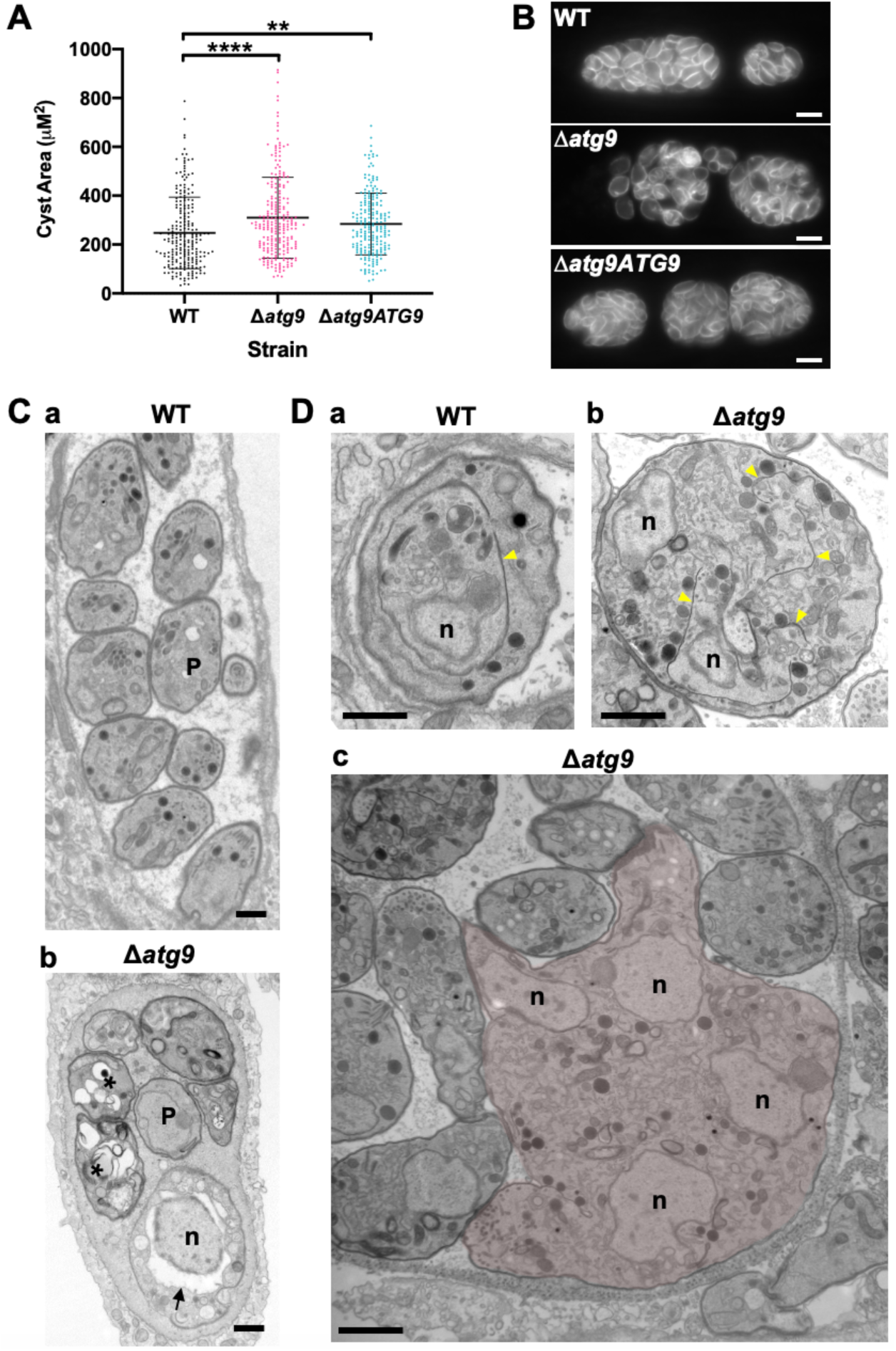
Canonical autophagy is required for normal bradyzoite morphology and cell division. (A) Quantification of cyst size based on staining for the cyst wall with *Dolichos* lectin. Each dot represents the size of one cyst. Data are merged from 3 biological replicates, each with a minimum of 52 cysts analyzed per sample type, per biological replicate and a minimum total of 199 cysts analyzed per sample type across 3 biological replicates. Bars indicate mean ± S.D. Statistical comparisons were done using a Kruskal-Wallis test with Dunn’s multiple comparisons. Significance is indicated **, p< 0.01; ****, p<0.0001. Non-significant differences are not indicated. (B) Immunofluorescence imaging of bradyzoites stained with anti-TgIMC1 revealed disorganization of the inner membrane complex in Δ*atg9* parasites and a “bloating” phenotype. Scale bar, 10 µm. (C) TEM showing a cyst containing normally developed bradyzoites from WT (a) and aberrant Δ*atg9* bradyzoites (b), with highly vacuolized, dying parasites (asterisks) or abnormally enlarged nuclear envelope (arrow). P, parasite. (D) TEM showing normal endodyogeny for WT bradyzoites with a nascent daughter containing well-organized organelles surrounded by the IMC (yellow arrowhead in panel a), and aberrant parasite division with poor packaging of organelles by the IMC (b), resulting in enlarged multinucleated Δ*atg9* bradyzoites (pseudocolorized parasite, c). Note that WT and mutant parasites are shown at the same magnification for direct comparison. All scales bars are 1 μm.

### A lack of canonical autophagy leads to ultrastructural abnormalities in the VAC of bradyzoites

The VAC performs a lysosome-like function in *T. gondii* tachyzoites and bradyzoites and is the site for proteolytic turnover of exogenously and endogenously derived proteinaceous material, including autophagosomes (9,10,27,28). Compared to the normal small and mixed electron density appearance of the VAC in WT bradyzoites (**Figure 5Aa**), TEM imaging revealed various abnormalities for this organelle in Δ*atg9* bradyzoites including some VACs that appeared enlarged and empty (**Figure 5Ab**). More commonly, other Δ*atg9* VACs were filled with highly electron dense material (**Figure 5Ac**), which sometimes included organellar remnants (**Figure 5Ad**). Whereas no WT (DMSO) bradyzoites had an electron dense VAC, 12.3% of Δ*atg9* bradyzoites showed this phenotype, suggesting a link between TgATG9 and proteolytic digestion in the VAC. The observation of electron dense material in Δ*atg9* bradyzoites (DMSO) is similar to, albeit much lower than LHVS-treated WT bradyzoites (**Figure 5Ba,b**), for which 93.1% showed an electron dense VAC. The VAC of LHVS-treated Δ*atg9* bradyzoites was sometimes only partially electron dense (**Figure 5Bc**), but the great majority (94.2%) of such parasites showed electron dense material (**Figure 5Bd**). This is consistent with our earlier observation that LHVS-treatment increases the size of dark puncta in Δ*atg9* bradyzoites, albeit not to the same degree as LHVS-treated WT or Δ*atg9ATG9* bradyzoites.

**Figure 5.**
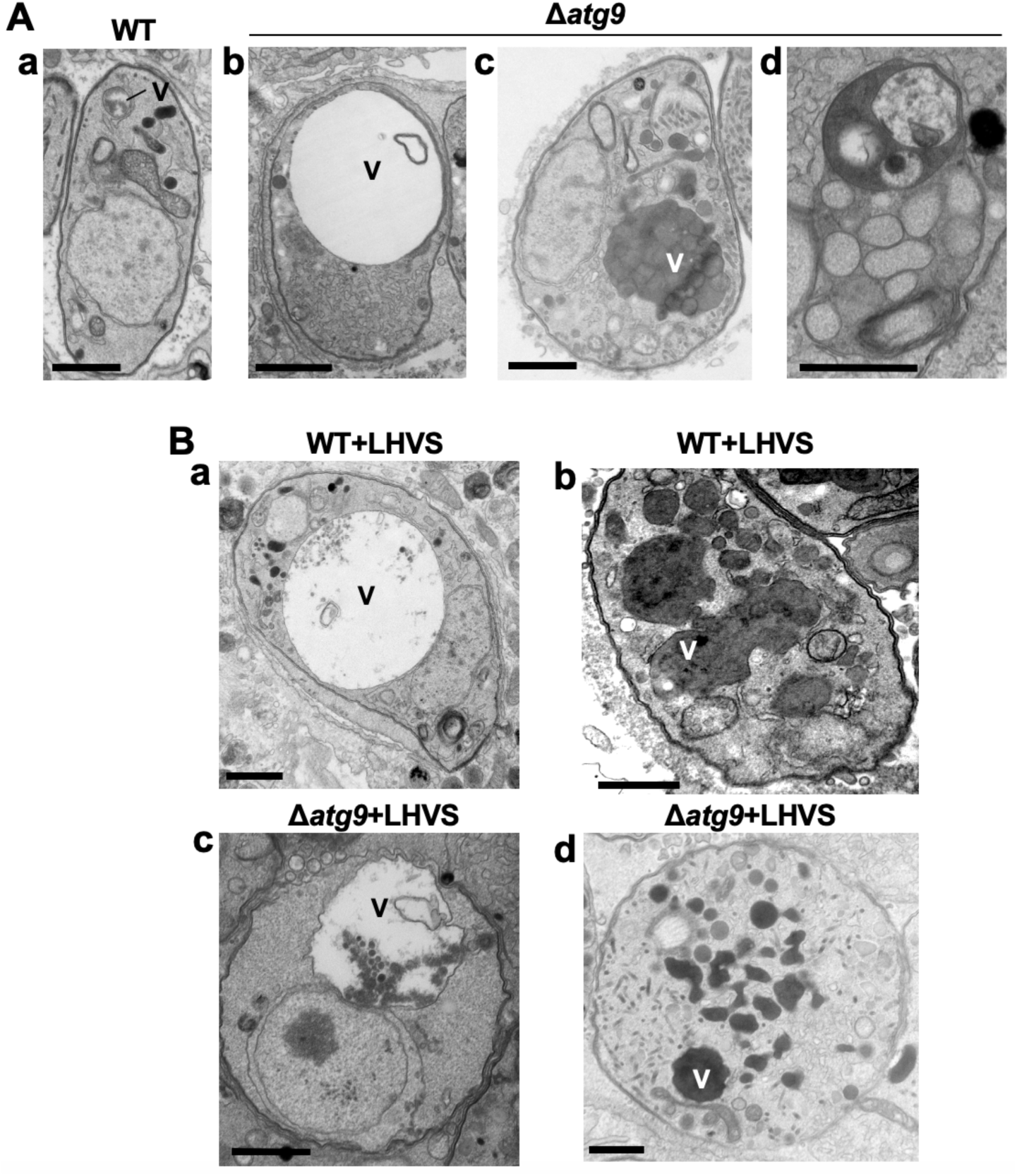
A lack of canonical autophagy leads to ultrastructural abnormalities in the VAC of bradyzoites. (A) TEM showing a normal VAC (V) in WT bradyzoites (panel a). Three types of VAC abnormalities were observed in Δ*atg9* bradyzoites: a very enlarged electron-lucent compartment (panel b), small electron-dense vesicles (panel c) or a very enlarged electron-dense compartment (panel d). (B) TEM of VAC in WT and Δ*atg9* bradyzoites treated with LHVS showing for both strains very large VAC with e-lucent content (panels a and c) or electron-dense content (panels b and d). All scales bars are 1 μm.

### Multimembrane structures and abnormal mitochondria in Δatg9 bradyzoites

Autophagic structures are typically transient and thus are rarely observed unless autophagic flux (the production and turnover of autophagosomes) is affected by increased biogenesis, reduced degradation, or both. This property is consistent with our inability to see autophagic structures by TEM in control treated WT (DMSO) bradyzoites, whereas 6.9% of WT bradyzoites treated with LHVS harbored such structures (**Table 1**). Interestingly, 6% of control treated and 10.1% of LHVS-treated Δ*atg9* bradyzoites also showed vesicular structures displaying two or more membranes, as exemplified in **Figure 6A**. Although it is uncertain whether these structures are autophagosomal, these observations suggest that TgATG9 is not required for the biogenesis of these multimembrane structures, and that ablation of TgATG9 potentially reduces the rate at which vesicular material is turned over by the VAC.

**Figure 6.**
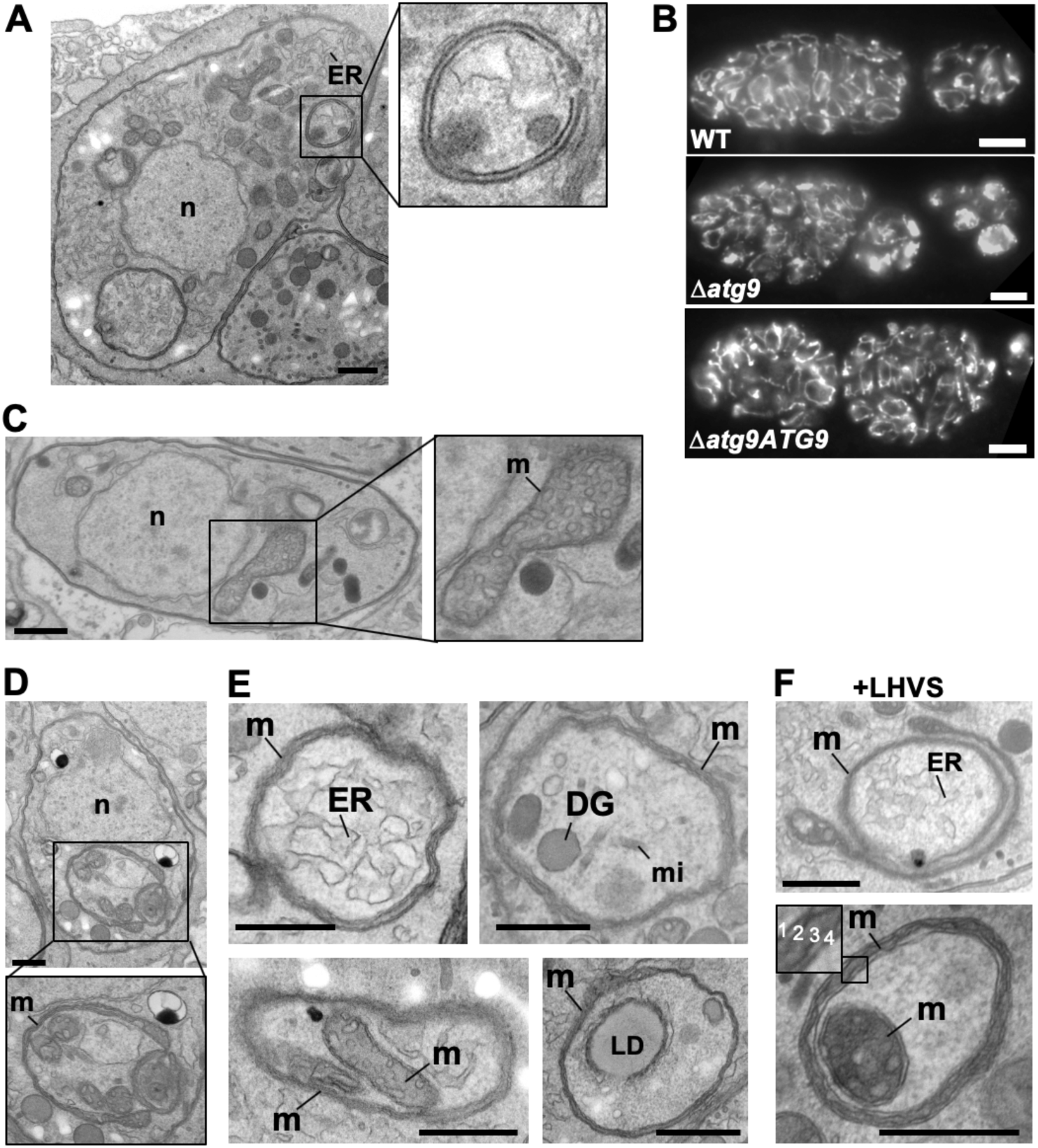
Autophagic structures and abnormal mitochondria in Δ*atg9* bradyzoites. (A) TEM of Δ*atg9* parasites showing a double membrane structure containing cytoplasmic components including ER. (B) Immunofluorescence imaging of bradyzoites stained with α-F_1_F_0_ ATPase revealed an aberrant mitochondrial network in some Δ*atg9* parasites. Scale bar, 10 µm. (C) TEM showing a normal mitochondrion in WT bradyzoites. (D) TEM of Δ*atg9* bradyzoites presenting thin mitochondria (m) in a horseshoe conformation with bulbous ends, wherein cristae can be seen. Nucleus (n) is denoted. (E) TEM of Δ*atg9* parasites revealing mitochondrial profiles (m) enveloping many organelles, including endoplasmic reticulum (ER), dense granules (DG), micronemes (mi), mitochondria (m) and lipid droplets (LD). (F) Examples of ER and mitochondrial section wrapped by the mitochondrion after bradyzoite treatment with LHVS. Inset of the lower panel illustrates the 4 membranes observed in such structures. All TEM scale bars are 500 nm.

In addition to the other abnormalities described above, the mitochondria of Δ*atg9* bradyzoites showed strikingly aberrant appearance. Whereas the mitochondria in WT bradyzoites displayed their typical tubular network, the mitochondria of Δ*atg9* bradyzoites appeared fragmented and punctate when observed by IFA (**Figure 6B**). Closer inspection by TEM confirmed that while the mitochondria of WT bradyzoites appeared normal, with internal cristae throughout (**Figure 6C**), 22.6% of Δ*atg9* bradyzoites had extremely thin mitochondria that presented a horseshoe-like appearance with bulbous ends, wherein cristae were present (**Figure 6D**). Often, the two ends of the U-shaped structure appeared to fuse together, encapsulating components of the cytoplasm, including ER, secretory organelles (dense granules and micronemes), lipid droplets and even other aspects of the mitochondria itself (**Figure 6E**). That such structures were comprised of 4 membranes (e.g., **Figure 4F** inset) clearly distinguished them from autophagosomes and the abnormal double-membrane vesicles observed. LHVS-treatment of Δ*atg9* bradyzoites did not seem to change the appearance or frequency (23.7%) of the abnormal mitochondria (**Figure 6F**). These thin mitochondria were never seen in control treated WT bradyzoites, but they were observed after LHVS-treatment at a frequency of 11.9%. Taken together, these findings suggest that impairment of autophagic flux due to ablation of TgATG9 or inhibition with LHVS leads to striking mitochondrial abnormalities in bradyzoites.

### Canonical autophagy is critical for bradyzoite viability and persistence

Observations by electron microscopy suggested that 30% of Δ*atg9* cysts showed signs of degeneration (**Table 1**), implying that a functional autophagy pathway is important for bradyzoite fitness. To more directly assess the role of TgATG9 in bradyzoite viability we differentiated WT, Δ*atg9* and Δ*atg9ATG9* for 7 days *in vitro* before treating them with vehicle or LHVS for 7 or 14 days under continued differentiation conditions and assessing viability by qPCR/plaque assay. Consistent with previous studies (28), LHVS treatment compromised bradyzoite viability, particularly after 14 days of treatment (**Figure 7A,B**). As expected, WT and Δ*atg9ATG9* bradyzoites treated with DMSO remained viable at both time points. However, vehicle treated Δ*atg9*-DMSO bradyzoites were much less viable than Δ*atg9ATG9* at both 7 and 14 days of treatment (corresponding to 14 and 21 days of differentiation). Δ*atg9* bradyzoites treated with LHVS showed a trend toward lower viability compared to LHVS treated WT and Δ*atg9ATG9*. We repeated the experiment to perform statistical comparisons of viability after 14 days of differentiation without treatment. The findings confirmed that Δ*atg9* bradyzoites are significantly less viable than WT or Δ*atg9ATG9* bradyzoites (**Figure 7C**). We also performed plaque assays on WT, Δ*atg9* and Δ*atg9ATG9* tachyzoites and found each strain formed plaques equally well, with no significant differences in the number of plaques formed between each strain (**Figure S2**). This indicates TgATG9 is non-essential in tachyzoites. It also suggests host cell invasion is not measurably impacted in parasites lacking TgATG9 (**Figure S2**) and that the lower plaque number recorded in Δ*atg9* bradyzoites is due to reduced parasite viability within the cyst, as opposed to impaired host cell invasion as parasites were placed onto a new monolayer for plaquing.

**Figure 7.**
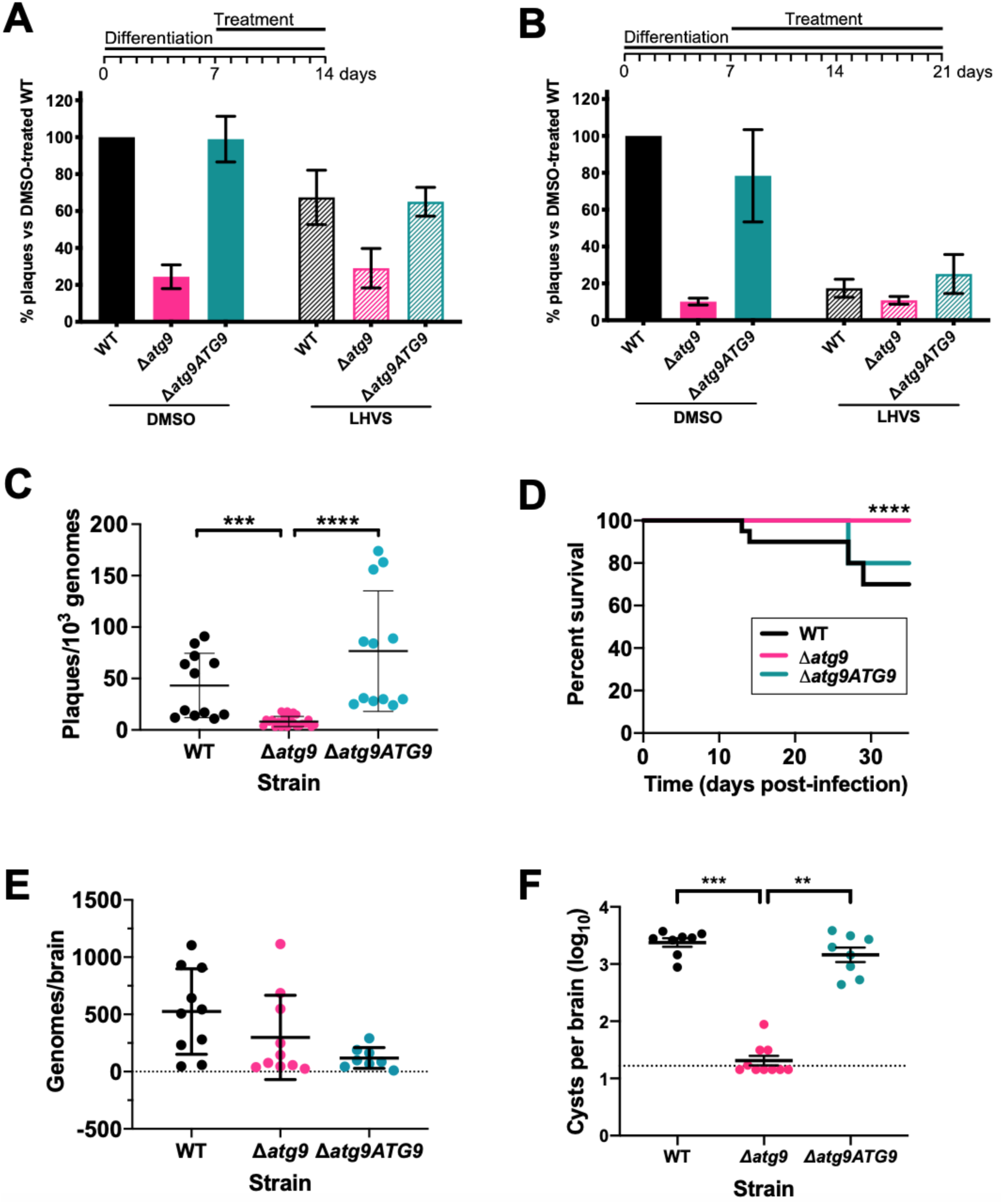
Autophagy is required for bradyzoite viability and the persistence of tissue cysts. Quantification of bradyzoite viability by qPCR/plaque assay after differentiation media for one week followed by treatment with DMSO (vehicle control, blue) or LHVS (red) for a further one week (A) or two weeks (B). Data are from 3 biological replicates each with 3 technical replicates and are normalized to WT set at 100%. Bars represent mean ± S.E.M. (C) Viability of untreated bradyzoites after 2 weeks of differentiation *in vitro*. Each dot represents viability from one technical replicate. Data are merged from 4 biological replicates. Statistical comparisons were done using a Kruskal-Wallis test with Dunn’s multiple comparisons. Significance is indicated as ***, p<0.001; **** p<0.0001. Non-significant differences are not indicated. (D) Analysis of virulence based on survival of infected mice. Mice were infected intraperitoneally with 150 tachyzoites of each strain. Infected mice that became moribund before the endpoint at day 35 post-infection were humanly euthanized. Mice infected with Δ*atg9* showed significantly better survival based on Mantel-Cox log-rank test (****, p<0.0001). (E) Analysis of parasite dissemination to the brain. Brains were harvested from infected mice at 7 days post-infection and the number of genomes present in each brain homogenate was determined by qPCR. No significant differences were seen between strains based on analysis with a Kruskal-Wallis test with Dunn’s multiple comparisons. (F) Analysis of brain cyst burden in infected mice. Brains were harvested from mice 5 weeks post-infection and cysts were counted in blinded samples of brain homogenates by microscopy. Data is plotted as log_10_ cysts per brain. The dotted line intersecting the Y axis represents the limit of detection (<33 cysts/brain). Samples for which no cysts were observed in the 30 µl of homogenate analyzed were given a value that is half the limit of detection. Bars represent mean ± S.E.M and statistical comparisons were done using a Kruskal-Wallis test with Dunn’s multiple comparisons. Significance indicated as **, p<0.01, ***, p<0.001. Non-significant differences are not indicated.

**Figure S2.**
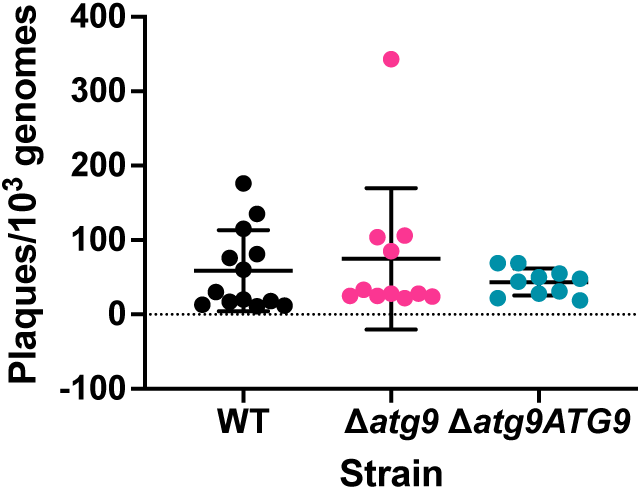
TgATG9 is not required for tachyzoite plaque formation. Tachyzoites were isolated and applied to fresh cell monolayers for plaque formation over the course of 10 days. Plaques were normalized to 10^3^ genomes of input parasites, as measured by qPCR. Each dot represents one technical replicate from 3 (Δ*atg9ATG9*) or 4 (WT and Δ*atg9*) biological replicates. No statistically significant differences were found using a Kruskal-Wallis test with Dunn’s multiple comparisons.

To determine whether canonical autophagy is necessary for the persistence of *T. gondii* tissue cysts *in vivo*, we infected mice with WT, Δ*atg9* and Δ*atg9ATG9* tachyzoites. Consistent with a previous study (31), we found that TgATG9 is necessary for normal virulence. Whereas 30% (6/20) and 20% (2/10) of mice infected with WT or Δ*atg9ATG9*, respectively, failed to survive to 35 days (5 weeks) post-infection, all mice (10/10) infected with the Δ*atg9* strain survived (**Figure 7D**). qPCR analysis of brain homogenate during the acute stage at day 7 post-infection showed that initial parasite brain burden did not differ significantly between mice infected with WT, Δ*atg9* and Δ*atg9ATG9* (**Figure 7E**), suggesting that TgATG9 deficiency does not affect parasite dissemination to the brain. By contrast, in mice that survived to 5 weeks post-infection we observed an ∼100-fold decrease in the brain cyst burden for mice infected with Δ*atg9* compared to those infected with WT or Δ*atg9ATG9* (**Figure 7F**). Taken together our findings suggest that canonical autophagy is critical for *T. gondii* bradyzoite long-term viability and persistence within tissue cysts *in vivo*.

## DISCUSSION

Autophagy is a complex intracellular pathway that facilitates the removal of endogenous proteins and organelles in eukaryotic cells. Often regarded as a starvation response to recycle amino acids for the production of new proteins, this process can also serve to remove misfolded proteins and damaged organelles, “reshape” the cell during development and, for mammalian cells, to remove pathogens (xenophagy) (18). In apicomplexan parasites, many autophagy-related proteins have been repurposed to have an additional role in maintaining the apicoplast, a remnant organelle derived from phagocytosis of an algal ancestor that is essential for parasite viability (22,24,29,32). Many of the autophagy-related proteins essential in this process, such as TgATG8, typically function “downstream” in the autophagy pathway (as depicted in **Figure 1B**). While a number of proteins involved in apicoplast maintenance have been studied and shown to be essential in *T. gondii* and *Plasmodium* spp. (22–24,29,32–35), little is known about the role of proteins that function earlier in the pathway. The autophagy-related protein TgATG9 has previously been shown to be non-essential for survival of *T. gondii* tachyzoites within the PV *in vitro* (31).

A cathepsin L enzyme in *T. gondii*, TgCPL, has been shown to be a major protease required for the turnover of exogenous and endogenous proteinaceous material in the lysosome-like organelle, termed the VAC (9,28). Similar to TgATG9, disruption of TgCPL has been found to be non-essential for tachyzoite growth and replication within the PV *in vitro* (5,15). This is likely due to parasites having sufficient access to nutrients (e.g., amino acids) from the host cell cytoplasm to support their rapid growth (9). Conversely, ablation of TgCPL function is catastrophic in chronic-stage bradyzoites, with TgCPL-deficient bradyzoites failing to survive *in vitro* and *in vivo* (27,28). Accumulation of autophagic material in the VAC of TgCPL-deficient bradyzoites coincided with loss of viability (27,28). This raised the question that although canonical autophagy had been found to be dispensable in intracellular tachyzoites, was it an important pathway in chronic stage bradyzoites? Here we demonstrated that removal of the *TgATG9* gene substantially reduces the accumulation of autophagic material in bradyzoites treated with LHVS to inhibit the enzymatic activity of TgCPL. Importantly, we also demonstrated that genetic ablation of *TgATG9* resulted in bradyzoite death *in vitro*, in both TgCPL-proficient and LHVS-treated (effectively TgCPL-deficient) parasites. When autophagic material was previously reported to accumulate in Δ*cpl T. gondii* bradyzoites (28), it was possible that autophagy was triggered in response to starvation, resulting from a general block in TgCPL-dependent protein turnover in the VAC. However, we found that the survival of untreated and DMSO-treated (TgCPL-proficient) Δ*atg9* bradyzoites was severely compromised *in vitro*, supporting the hypothesis that autophagy is a primary pathway in chronic-stage *T. gondii* which is essential for survival within the cyst. This is supported by *in vivo* experiments performed in mice in which Δ*atg9* brain cysts were severely reduced at 5-weeks post-infection. Having detected Δ*atg9* parasites in the brains of mice 1-week post-infection, we stipulate that autophagy is a survival mechanism in *T. gondii* that facilitates parasite persistence within the cyst.

Upon closer inspection of *in vitro* bradyzoites by TEM, we noticed that treatment with LHVS led to numerous ultrastructural abnormalities in the VAC in both WT and Δ*atg9*. For example, enlarged “empty” VACs were observed following LHVS treatment, which were also detected in Δ*atg9* bradyzoites treated with DMSO. Taken together, this suggests that TgATG9 and possibly autophagic flux play an important role in VAC homeostasis. As mentioned above, inhibition of TgCPL with LHVS predominantly results in enlargement of dark puncta, which correspond to accumulation of electron-dense and autophagic material in the VAC, shown here and previously documented by us (28). We observed a less pronounced enlargement of dark puncta in LHVS-treated Δ*atg9* bradyzoites compared with LHVS-treated WT or Δ*atg9ATG9*. This, coupled with our observation that 90% of LHVS-treated *Δatg9* bradyzoites have an electron-dense VAC, suggests that some trafficking of proteinaceous material to the VAC still occurs in such parasites. However, since we did not see an increase in the number of CytoID-positive puncta in LHVS-treated Δ*atg9* bradyzoites compared with DMSO-treated Δ*atg9*, this proteinaceous material does not seem to be derived from canonical autophagosomes. We propose two possibilities, that bradyzoites uptake host proteins that are trafficked to the VAC for degradation via the endolysosomal system or that, in the absence of TgATG9, bradyzoites have an alternative mechanism of autophagy. In the first possibility, although uptake of host-derived proteins across the cyst wall has not been directly shown in bradyzoites, it has been established in acute stage tachyzoites, whereby fluorescent host protein has been found to accumulate in the VAC of TgCPL-deficient parasites (9,10,27,36). In the second possibility, as discussed further below, we observed the formation of vesicles with two or more membranes in *Δatg9* bradyzoites by TEM microscopy and while we are currently unable to determine the nature of these vesicles, it remains possible that they represent alternative autophagosome-like structures that do not stain well with the cationic amphiphilic dye CytoID. In either case, the inability of Δ*atg9* bradyzoites to survive and persist suggests that neither of these alternative pathways can rescue the function of canonical autophagy in *T. gondii* bradyzoites.

TEM imaging also identified numerous structural anomalies that were present in Δ*atg9* bradyzoites but absent in the parental strain. Structural abnormalities included bradyzoites with a division defect, manifested by, for example, the observation of four nuclei within a single bradyzoite. Although endopolygeny has previously been reported in *T. gondii* bradyzoites (37), we found it to occur in less than 1% of WT parasites. However, a sharp increase in the number of mother cells containing multiple daughter nuclei was observed in the *TgATG9* knockout strain, with approximately 30% of bradyzoites showing this phenotype. This finding implicates TgATG9 in bradyzoite replication and development, although whether the protein has a direct role or if this observation was an indirect consequence from a lack of canonical autophagy is unclear. This is notably in contrast to what has previously been observed in TgATG9-deficient *T. gondii* tachyzoites, in which the parasite lytic cycle (acute stage invasion, replication and egress) was found to be unaffected by the gene knockout (31). Other structural abnormalities observed in Δ*atg9* bradyzoites included a disorganization of the inner membrane complex and the occurrence of vesicular objects that had multiple membranes. In some cases, these structures had two double membranes and could represent autophagosome-like vesicles produced in a TgATG9-independent manner. Although ATG9 is required for efficient autophagy, alternative vesicle and puncta formation has been found to occur in other eukaryotic organisms in which *ATG9* has been knocked out or silenced (38–40).

However, we also observed structures with at least four membranes that appeared to be derived from the mitochondria, with cristae clearly visible. A primary role of ATG9 in autophagy is to initiate the formation of the lipid bilayer for the developing phagophore (41,42). Although ATG9 has been shown to shuttle between the surface of mitochondria to the phagophore assembly site in yeast (43,44), the Golgi apparatus has been shown to be the major source of ATG9 pools, particularly in other eukaryotes (45–47). In a previous study, TgATG9 was not found to co-localize with mitochondria in tachyzoites, but did co-localize with the Golgi and other organelles including the VAC, suggesting these as potential sources of TgATG9 in this parasite (31). Ablation of TgATG9 was not found to have an extensive impact on mitochondrial morphology in tachyzoites, but it was not assessed at the ultrastructural level (31). Here, our TEM findings of TgATG9-deficient bradyzoites frequently showed multiple fragmented and elongated mitochondria within a single parasite. An increase in mitochondrial elongation and fragmentation was shown to occur in *Drosophila* deficient in the ATG2/ATG18/ATG9 complex (48). Most intriguingly the mitochondrial elongations we observed in TgATG9-deficient *T. gondii* bradyzoites seemed to form mitochondrial vacuoles strikingly similar to “m-vacuoles” previously reported in the protist, *Dictyostelium* (49,50), whereby the mitochondria has been shown to elongate with bulbous ends and adopts a horseshoe conformation that encapsulates the cytoplasm within differentiating prespore cells. Spherical mitochondrial structures have also been observed in mammalian cells treated with CCCP (51,52), which collapses the mitochondrial membrane potential. These spherical structures were reported to encapsulate a variety of cytoplasmic structures and organelles, which is similar to the apparent sequestration of ER, secretory organelles, lipid droplets, and even parts of the mitochondria itself in Δ*atg9* bradyzoites. It is worth noting that previous reports have implicated ATG8 in organelle clearance in *Plasmodium* spp, which is essential to parasite development (34,53). We speculate that the encapsulation of organelles by multimembrane vesicles (including apparent m-vacuoles) observed here could represent an attempt at organelle clearance in the absence of ATG9-dependent canonical autophagy in the related apicomplexan *T. gondii*. However, the extent to which the abnormal mitochondria are a consequence of or a response to deficient autophagy remains to be determined.

Expectedly, most conservation of *T. gondii* autophagy-related proteins was found in the closely related Sarcocystidia *Hammondia hammondii, Neospora caninum* and *Sarcocystis neurona.* While we identified matches across apicomplexans to many of the *T. gondii* autophagy-related proteins that are involved in maintenance of the apicoplast, little to no conservation was identified in parasites outside of the coccidia for apicoplast-independent canonical autophagy proteins (e.g., ATG1, ATG2, ATG9 and Prop1). The potential conservation of canonical autophagy in coccidia supports the hypothesis that this pathway promotes survival within the cysts that persist either in tissues or in the environment. Although we did not identify canonical autophagy genes in Cryptosporidia, which also use environmentally resilient oocysts for fecal-oral transmission, such parasites are known to have a severely reduced genome and thus might have evolved other strategies for long-term survival. Not only did we identify matches to *T. gondii* ATG1, ATG9 and TgProp1 in *Chromera velia* and *Vitrella brassicaformis*, but we also found that many of the apicoplast-related autophagy proteins had better matches to proteins in these non-parasitic, photosynthetic protists than they did to other apicomplexan parasites. *C. velia* and *V. brassicaformis* were originally identified as endosymbionts of Scleractinia (stony corals) and represent some of the closest known non-parasitic relatives of the Apicomplexa (54–56). Although it is not clear what role autophagy plays in their life cycle, matches for canonical autophagy proteins between these endosymbiotic relatives and certain apicomplexans indicates canonical autophagy was conserved in a number of cyst-forming apicomplexans from a common ancestor and was lost in several apicomplexans that do not form tissue cysts (e.g. *Cryptosporidium spp, Plasmodium spp*, and *Babesia spp*).

Although our findings suggest a critical role for TgATG9 in canonical autophagy and persistence, virtually nothing is known about the initiation and biogenesis of autophagic structures in *T. gondii*. Gaining such insight will require the identification and interrogation of other autophagy-related proteins that function early in the pathway including the TgATG1 and TgATG2 homologues. Identifying other, potentially divergent, components of the early pathway will permit their evaluation as candidates for selectively disrupting the pathway to curb chronic infection.

## MATERIALS AND METHODS

### Ortholog identification

We sought to identify putative orthologs of *T. gondii* autophagy-related proteins across a number of closely related apicomplexan parasites (*Hammondia hammondi, Neospora caninum, Sarcocystis neurona, Eimeria spp, Cyclospora cayetanensis, Plasmodium falciparum, Babesia spp* and *Cryptosporidium parvum*) and distantly related photosynthetic, non-pathogenic relatives (*Chromera velia* and *Vitrella brassicaformis*). Amino acid sequences for *T. gondii* autophagy-related proteins were retrieved from ToxoDB.org and BLASTed against online databases for the relevant species listed above (ToxoDB.org, PlasmoDB.org, PiroplasmaDB.org and CryptoDB.org). Sequence hits with E values of <10^−3^ were considered a putative match. Negative logarithms of the E values were plotted into a heat map to demonstrate conservation of autophagy-specific proteins in cyst-forming protozoans. To perform a negative logarithm transformation on highly conserved hits that returned E values of 0.0, these E values were set at 10^−200^.

### Generation of the S/ATG8 T. gondii strain

The SAG1 promoter cassette was amplified using primers ML2475/ML2476 containing SpeI and XbaI restrictions sites, respectively, and cloned into the DHFR-TetO7 vector (Morlon-Guyot et al. Cell Microbiol. 2014) to yield the DHFR-pSAG1 plasmid. Then, using this plasmid as a template, the DHFR-pSAG1 cassette was amplified by PCR with primers ML2669/ML2670 and cloned downstream of the GFP-coding cassette in the NsiI-digested pGFP-TgAtg8 plasmid (35). Clones were verified for correct insert orientation by sequencing. Using this plasmid as a template, a 3,706 bp fragment containing the DHFR expression cassette, the SAG1 promoter and the GFP-coding fragment was amplified by PCR using the KOD polymerase (Merck) and the ML2477/ML2664 primers that included regions for integration by double homologous recombination. A protospacer sequence targeting the native TgATG8 promoter was generated by annealing primers ML2885/ML2886 and inserting the corresponding fragment at the BsaI site of vector pU6-Cas9 (30). This plasmid was co-transfected with the DHFR-pSAG1-GFP donor sequence in tachyzoites of the Prugniaud strain for integration by CRISPR/CAS9, and transgenic parasites were then selected with pyrimethamine and cloned by limiting dilution.

### Generation of the TgATG9 knock out cell line

The HXGPRT cassette was amplified from the pmini-HXGPRT plasmid (57) using primers ML2465/ML2466 containing the HindIII and BamHI restriction sites, respectively. This cassette was then inserted into the HindIII-BamHI digested pTub5/CAT-TgAtg9 plasmid, which was previously generated for creating a TgATG9 knock-out in a RH strain and thus already contained 5’ and 3’ TgATG9 homology regions for gene replacement (Nguyen et al., Cell Microbiol. 2016). The resulting HXGPRT-TgATG9 plasmid was then linearized with KpnI/NotI. A protospacer sequence targeting the 5’ of the TgATG9 coding sequence was generated by annealing primers ML2467/ML2468 and inserting the corresponding fragment at the BsaI site of vector pU6-Cas9 (Sidik et al., Cell, 2016). The linearized HXGPRT-TgATG9 plasmid, serving as a donor sequence, and the TgATG9-targeting Cas9-expressing plasmid were co-transfected in ΔHXGPRT Prugniaud tachyzoites. Transgenic parasites were then selected with mycophenolic acid and xanthine, and subsequently cloned by limit dilution.

### Genetic complementation

The coding sequence of the *TgATG9* gene was amplified from a *T. gondii* cDNA library (derived from the ME49 strain) and 1,500 bp of the *TgATG9* promoter was amplified from genomic DNA of the PruΔ*hxg* strain. The amplified *TgATG9* and its promoter region were ligated to one another and subsequently inserted into a complementation plasmid used in a previous study by Di Cristina *et al* (28) for complementing the *T. gondii TgCPL* gene. The *TgCPL* coding sequence in the plasmid was removed by restriction digest and the *TgATG9* expression cassette was inserted by ligation. The plasmid was then linearized by digestion with PsiI and used to transfect 1 x 10^7^ Δ*atg9* tachyzoites. The linearized plasmid was incorporated into the genome of Δ*atg9* tachyzoites by random integration. Based on the presence of a resistance cassette in the transfection plasmid, positively transfected parasites were selected using bleomycin prior to isolating clones by limiting dilution. Clonal populations were screened for TgATG9 expression by probing fixed parasites with α-HA antibodies.

### Bradyzoite Differentiation

For all bradyzoite conversion, tachyzoites were mechanically lysed by scraping infected HFF monolayers that were then passed sequentially through 20G and 23G syringes and a 3 μm filter. Filtered parasites were then counted and allowed to infect fresh monolayers of HFF cells for 24 h. Bradyzoite differentiation was induced using alkaline pH medium and ambient CO_2_ (58–60). Briefly, 24 h after parasites were applied to HFF monolayers, DMEM media was replaced for an alkaline differentiation media (RPMI without NaHCO_3_, 50 mM HEPES, pen/strep, 1% FBS, pH 8.25). Differentiation media was replaced daily.

### Immunoblotting

Tachyzoites were harvested while they were still largely intracellular. Bradyzoites were generated by induced differentiation with 1% FBS for 7 days in ambient CO_2_ condition. Parasites were lysed in RIPA buffer for 10 min at 4°C with shaking and pelleted by centrifuging at 10,000 *g* for 10 min at 4°C. Supernatants were collected and mixed with sample buffer and β-mercaptoethanol. Protein lysates from approximately 2.5 x10^6^ parasites were subjected to electrophoresis on 7.5% SDS-PAGE gels and transferred onto 0.45 µm nitrocellulose membranes (Bio-Rad, cat. # 1620115). Buffer containing a high concentration of both Tris and glycine (50 mM Tris, 380 mM glycine, 0.1% SDS, and 20% methanol) were used for transferring proteins to the membrane over the course of 10 h at 20 V at 4°C. Following transfer, the membrane was blocked with 5% milk in PBS (with 0.05% Tween X-114) for 30 min at room temperature. Primary antibodies were diluted in wash buffer (1% milk in PBS with 0.05% Tween 20) and applied to membranes overnight at 4°C. The primary antibodies used include mouse anti-HA (1:2500; clone 16B12, Biolegend, cat. # 901501), mouse anti-GFP (1:500; clones 7.1 and 13.1, Roche, cat. # 11814460001), rabbit anti-TgATG8 (1:500; (35)), mouse anti-α-tubulin (1:2,000; clone B-5-1-2, Sigma-Aldrich, cat. # T5168) and mouse anti-MIC2 (1:2500; 6D10). Membranes were washed three times with wash buffer before incubation with HRP-conjugated secondary antibodies (1:2500) for 1 h at room temperature. Proteins were detected using SuperSignal West Pico PLUS Chemiluminescent Substrate (Thermo Fisher Scientific, cat. #1863096) or SuperSignal West Femto (Thermo Fisher Scientific, cat. #34094). The Syngene Pxi imaging system was used to detect signals.

### Staining autophagic material with CytoID

Tachyzoites were differentiated to bradyzoites for 7 days and then treated with either 1 µM LHVS or an equal volume of DMSO for 1 day. The CytoID™ Autophagy Detection Kit 2.0 (Enzo) was used to stain autophagosomes within live bradyzoites prior to fixation with 4% paraformaldehyde, following the manufacturer’s instructions, and the cyst wall was stained using Rhodamine labelled *Dolichos biflorus* agglutinin (1:400; Vector Laboratories, cat. # RL-1032). The number and size of CytoID positive structures were measured using CellProfiler (61). The boundaries of each cyst were identified manually based on *Dolichos* staining. Raw images for CytoID staining were first corrected for uneven illumination/lighting/shading to reduce uneven background signal. CytoID signal was identified using Otsu two-classes thresholding method. The image processing pipeline is available at: https://cellprofiler.org/examples/published_pipelines. Measurements of cytoID puncta were done automatically within CellProfiler. The definition of all the measurements can be found at: http://cellprofiler-manual.s3.amazonaws.com/CellProfiler-3.0.0/modules/measurement.html.

For probing autophagic material in S/Atg8 bradyzoites, tachyzoites were converted for 7 days. After this time LHVS (1 µM) was added for either 1 or 7 days. An equal volume of DMSO was added as a vehicle control treatment. CytoID was then performed as described above.

### Immunofluorescence

For immunofluorescence assays (IFA), HFF monolayers were grown overnight on coverslips then infected with parasites for 1 h before they were washed twice with 37°C PBS, fixed for 20 min with 4% (w/v) paraformaldehyde in PBS, permeabilized with 0.1% Triton X-100 in PBS for 15 min, and blocked with 0.1% (w/v) fatty-acid free BSA (Sigma-Aldrich, cat. # 9048-46-8) in PBS. Rat anti-HA (Sigma-Aldrich, cat. # 11867423001) (1:500), mouse anti-GFP (1:500; clones 7.1 and 13.1, Roche, cat. # 11814460001), mouse MAb 45.56 anti-TgIMC1 (1:500; Gary Ward, University of Vermont), with mouse MAb anti-F_1_F_0_ATPase (1:2000; P. Bradley, UCLA) and rabbit anti-TgCPN60 (1:2,000; (62)) were used for 1 h of primary antibody staining in wash buffer consisting of 1% cosmic calf serum in PBS. Cover slips were rinsed three times and then washed three times for 5 min in wash buffer. Secondary antibodies were: goat anti-rat Alexa Fluor 594 (Invitrogen, cat. # A11007) (1:1000), goat anti-rabbit Alexa Fluor 594 (Invitrogen, cat. # A11012), goat anti-rabbit Alexa Fluor 488 (Invitrogen, cat. # A11008) and goat anti-streptavidin Alexa Fluor 350 (1:1000). They were used for 1 h diluted in wash buffer at 1:1000. DAPI (Sigma, cat. # D9542) was used at 1:200. Fluorescein and biotinylated *Dolichos biflorus* agglutinin were used at 1:400 (Vector Laboratories, cat. # FL-1031 and B-1035). Cover slips were then washed three times for 5 min in wash buffer and subsequently mounted on slides with 8 μl of prolong glass (Invitrogen, cat. # P36984) or mowiol. Images were taken on a Zeiss Axio Observer Z1 inverted microscope at 100X and analyzed using Zen 3.0 blue edition software.

### Bradyzoite Puncta (Fig 5A/B)

The accumulation of dense material within *in vitro* bradyzoite cysts was quantitatively measured using the puncta quantification assay, previously described (28). Briefly, HFF cell monolayers were grown on cover slips and infected with *T. gondii* tachyzoites. Twenty-four hours post-invasion, intracellular tachyzoites were differentiated to bradyzoites over the course of 7 days (as described above) and subsequently treated with 1 µM LHVS or 0.1% DMSO (vehicle control) for 24 hours. For direct immunofluorescence of bradyzoite cyst walls, *Dolichos biflorus* agglutinin conjugated with fluorescein or rhodamine (Vector labs, cat. #s FL-1031 and RL-1032-2, respectively) was diluted 1:400 and incubated with infected monolayers that had been fixed with 4% paraformaldehyde and made permeable with triton X-100. The fluorescent signal derived from DBA staining allows automatic detection of the bradyzoite-containing cyst area using ImageJ software. Cyst images were captured at 100X or 63X oil objective (3 biological replicates at each objective) as described above. Automatic bradyzoite cyst identification and puncta quantification was performed in ImageJ, using the following parameters described previously by Di Cristina *et al* (28). Max Entropy thresholding on the GFP channel was used to identify cysts. This was followed by the identification of objects with areas between 130 and 1,900 μm^2^. Particles (puncta) under the GFP mask (therefore within a cyst) were analyzed by automatic local thresholding on the phase image using the Phansalkar method, with the following parameters: radius = 5.00 μm; k value = 0.5; r value = 0.5. Puncta were measured from the resulting binary mask by particle analysis according to the following: size = 0.3–5.00 μm; circularity = 0.50–1.00.

### Transmission electron microscopy of in vitro bradyzoite cysts

Sample preparation for transmission electron microscopy was carried out as described by Di Cristina *et al* (28). Briefly, HFF monolayers grown in 6-well plates were infected with 5 × 10^4^ tachyzoites in D10 media. After O/N incubation at 37°C, 5% CO_2_, the D10 medium was replaced with alkaline media (pH 8.2) to induce bradyzoite conversion. Alkaline media was changed daily over the course of seven days, replacing with fresh alkaline media and plates were incubated at 37°C with 0% CO_2_. TEM preparation of samples was carried out by washing the infected monolayers with cold PBS three times, followed by fixation with 2.5% glutaraldehyde (EMS, cat. # 16210) in 0.1 M sodium cacodylate buffer (pH 7.4) for 1 h at RT. Fixed infected monolayers then were gently lifted using a cell scraper to detach large sheets and transferred to microcentrifuge tubes. Samples were centrifuged at 1,500g for 10 min at RT and washed three times with 0.1 M sodium cacodylate buffer (pH 7.4). Samples were stored in the same buffer at 4°C until processed for TEM as described in Coppens and Joiner (63) before examination with a Philips CM120 electron microscope under 80 kV.

### Bradyzoite Viability

Bradyzoite viability was assessed by combining plaque assay and quantitative polymerase chain reaction (qPCR) analysis of genome number, as previously described by Di Cristina *et al* (28). Briefly, HFF cell monolayers were infected with *T. gondii* tachyzoites in 6-well plates. Following host cell invasion tachyzoites underwent differentiation to bradyzoites as described above, resulting in the generation of *in vitro* tissue cysts. Differentiation was carried out over the course of 7 days, replacing the alkaline media daily. Following differentiation, parasites were treated with 1 µM LHVS or 0.1% DMSO (vehicle control) in differentiation media. Treatment was replaced daily for 7 and 14 days, respectively. Following the treatment period, the culture media in each well was replaced with 2 mL Hanks Balanced Salt Solution and cysts were liberated from the infected HFF monolayers by mechanical extrusion, by lifting cells with a cell scraper and syringing several times through 25-gauge needles. Then, 2 mL of pre-warmed 2× pepsin solution (0.026% pepsin in 170 mM NaCl and 60 mM HCl, final concentration) was added to each sample and samples left to incubate at 37°C for 30 min. Reactions were stopped by adding 94 mM Na_2_CO_3_, removing the supernatant after centrifugation at 1,500g for 10 min at RT and resuspending pepsin-treated parasites in 1 mL of DMEM without serum. Parasites were enumerated and 1,500 parasites per well were added to 6-well plates containing confluent monolayers of HFFs in D10 media, in triplicate. To allow for the formation of bradyzoite-derived plaques, parasites were left to grow undisturbed for 12 days. After this period, the number of plaques in each well was determined by counting plaques with the use of a light microscope. Five hundred μL of the initial 1 mL of pepsin-treated parasites was used for genomic DNA purification, performed using the DNeasy Blood & Tissue Kit (Qiagen). Genomic DNA was eluted in a final volume of 200 µL. To determine the number of parasite genomes per microliter, 10 µL of each gDNA sample was analyzed by qPCR, in duplicate, using the tubulin primers TUB2.RT.F and TUB2.RT.R (36). Quantitative PCR was performed using Brilliant II SYBR Green QPCR Master Mix (Agilent) and a Stratagene Mx3000PQ-PCR machine. The number of plaques that formed was then normalized to the calculated number of genomes present in the inoculating sample. Despite this normalization, these experiments can have considerable inter-experiment variation, which in this case necessitated secondary normalization to WT-DMSO within each biological replicate. Because secondary normalization to WT-DMSO precluded statistical comparison of this group to the others, we repeated the experiment and assessed viability after 14 days of differentiation without treatment.

### Mouse infection experiments

Seven-week-old female CBA/J mice (Jackson) were randomly assigned to groups and infected intraperitoneally (i.p.) with 150 tachyzoites of WT, Δ*atg9* or Δ*atg9ATG9* strains. Group sizes in each experiment are as follows: In the first experiment, 10 mice were infected with one of the three strains, 5 mice per strain were humanely euthanized 1-week post-infection to assess brain tachyzoite burden, 5 mice per strain were humanely euthanized 5-weeks post-infection to assess brain cyst burden. The second experiment was set up the same as the first experiment, however an additional 10 mice were also infected with WT and their survival assessed. Assessing the statistical differences in parasite burden between mice infected with each strain was complicated by the absence of data from the mice that did not survive infection. Because mice that die from the infection often have higher burden and become moribund and must be euthanized before the 5-week time point when cysts are counted, enumeration of cysts from those that survive probably underestimates differences. Data were pooled from the two independent experiments. The mouse brains were placed in a set volume of sterile PBS that provided a concentration of 500 mg of mouse brain per ml. Mouse brains were individually minced with scissors, vortexed and homogenized by three or four passages through a 21G syringe needle. Three 10 µl samples (30 µl total) of brain homogenates per infected mouse were analyzed by phase-contrast microscopy to enumerate cysts and the number of cysts per brain determined (scaled appropriately to the total volume of brain homogenate for that mouse). Samples for which no cysts were observed in the 30 µl of homogenate were given a value (17 cysts/brain) that is half the limit of detection (<33 cysts/brain). Mouse sample sizes were chosen based on previous studies. Mice were randomly assigned to groups and samples were blinded for enumeration of cysts. Animal studies described here adhere to a protocol approved by the Committee on the Use and Care of Animals of the University of Michigan.

Genomic DNA was extracted from 25 mg of brain homogenate tissue using the DNeasy Blood and Tissue Extraction kit (Qiagen) following the manufacturer instructions. SYBR green qPCR was performed on gDNA using 300 nM Tox9 and Tox11 primers and reaction conditions previously described (36).

### Tachyzoite plaque assays

Intracellular tachyzoites were purified from HFFs following standard protocols. Parasites were added to and left undisturbed on a monolayer of HFFs for 10 days, after which time plaques were counted. An aliquot of purified parasites was collected for gDNA extraction to normalize tachyzoite plaque numbers. Briefly, gDNA was extracted using the Phire Tissue Direct PCR Master Kit (Thermo Scientific; (64)). SYBR Green qPCR was performed for alpha-Tubulin following conditions previously used (36).

### Statistical analysis

Data were analyzed after removing outliers using ROUT with a Q value of 0.1% and testing for normality and equal variance. Since all data failed either test, a Kruskal-Wallis with Dunn’s Multiple Comparisons was used to compare groups within the same genotype or treatment. Mouse survival was analyzed using a Mantel-Cox log-rank test.

## Acknowledgements

We thank Christian McDonald and all members of the Carruthers lab for providing feedback on the study. The authors appreciate My-Hang (Mae) Huynh’s help with proofreading the manuscript. We also thank Mack Reynolds for help with CellProfiler, Aric Schultz for help with ImageJ, Michael Delannoy and other staff at the Johns Hopkins University Microscopy facility, and Drs. Peter Bradley, Boris Striepen, and Gary Ward for generously providing antibodies for this study. This work was supported by National Institutes of Health grants R01AI120627 (VBC and MDC) and R01AI060767 (IC) together with support from the Agence Nationale de la Recherche (ANR-19-CE15-0023) the Fondation pour la Recherche Médicale (FRM EQ20170336725) (SB) and the University of Perugia Fondo Ricerca diBase 2019 program of the Department of Chemistry, Biology and Biotechnology (MDC).

